# A Variational Autoencoder Model for Clustering of Cell Nuclei on Microgroove Substrates: Potential for Disease Diagnosis

**DOI:** 10.1101/2024.12.27.630482

**Authors:** Bettina Roellinger, Francois Thenier, Claire Leclech, Catherine Coirault, Elsa Angelini, Abdul I. Barakat

## Abstract

Various diseases including laminopathies and certain types of cancer are associated with abnormal nuclear mechanical properties that influence cellular and nuclear deformations in complex environments. Recently, microgroove substrates designed to mimic the anisotropic topography of the basement membrane have been shown to induce significant 3D nuclear deformations in various adherent cell types. Importantly, these deformations are different in myoblast cells derived from laminopathy patients from those in cells derived from normal individuals. Here we assess the ability of a variational autoencoder (VAE) and a Gaussian Mixture Model (GMM) to cluster patches of nuclei of both wildtype myoblast cells and myoblast cells with laminopathy-associated mutations cultured on microgroove substrates, and we explore the impact of image processing parameters on clustering performance. We show that a standard VAE with GMM is able to cluster nuclei based on their morphologies and degrees of deformations and that these clusters correspond to either wildtype myoblasts or myoblasts with LMNA mutations. The current results suggest that combining deep learning techniques with microgroove substrates enables automatic classification of nuclear deformations and thus provides a promising approach for easy and rapid diagnosis of pathologies that involve abnormalities in nuclear deformation.

## Introduction

A central question in biology that is pertinent at multiple scales is the relationship between form and function^1^. For instance, abnormalities in cell shape have long been recognized as hallmarks of cancer^2^, and nuclear morphometric parameters are routinely used by diagnostic pathologists to distinguish between benign and malignant cells^3^,^4^,^5^. There is now extensive evidence that overall cell shape and the organization of various intracellular organelles are exquisitely sensitive to the extracellular mechanical environment^6^,^7^,^8^,^9^. A prominent intracellular mechano-responsive organelle is the nucleus which exhibits alterations in position as well as changes in gene and protein expression in response to mechanical stimulation^10^,^11^. In the nucleus, mechanical stress resistance is thought to be provided principally by the nuclear lamina which acts as a stiff and load-bearing structure^12^. The lamina is a dense fibrillar network that underlies the inner nuclear membrane and consists of A-type (lamin A and C) or B-type (lamin B1 and B2) lamins and a number of lamin-associated proteins. The importance of the nuclear lamina in biological processes is underscored by the premature death of mice that lack functional lamin A/C^13^,^14^ and by the fact that mutations in the lamin A/C gene give rise to a complex set of pathological conditions collectively termed “laminopathies”. These disease phenotypes include cardiac and skeletal myopathies, lipodystrophies, and premature aging (Hutchinson-Gilford progeria syndrome). Several studies have correlated lamin A/C deficiency with misshapen nuclei. For instance, distorted nuclear shape has been demonstrated in fibroblasts from lipodystrophic patients with heterozygous R482Q/W mutations in the lamin A/C gene and in cells from C. elegans with reduced lamin levels^15^,^16^. In several experiments using multiple breast epithelial cell preparations, lamin A/C down-regulation consistently resulted in 30 to 70% of the cells having larger nuclei with atypical morphology^17^. These findings and others suggest that lamin expression and organization play a critical role in regulating nuclear size and shape.

Micro- and nano-scale surface patterning has recently emerged as a powerful and robust tool for non-invasive control of both cell shape and intracellular organization^18^. Planar adhesive line patterns and microstructured topographic surfaces of appropriate dimensions have been shown to induce nuclear elongation and alignment in several cell types^19^. We have recently demonstrated that the nuclei of human umbilical vein endothelial cells and human muscle precursor cells (myoblast) cultured on substrates consisting of arrays of microgrooves that mimic the anisotropic topography of the basement membrane underlying adherent cells deform significantly and assume different morphologies,^20^. In some cases, these deformations can lead to the nuclei being entirely trapped or “caged” within the microgrooves. This process of nuclear caging is transient and is associated with significant changes in nuclear mechanical properties^20^. Interestingly, the extent of nuclear deformations on microgroove substrates in normal myoblast cells appears to be significantly different from that in myoblast cells derived from laminopathy patients^20^. These results suggest that monitoring nuclear deformation on microgroove substrates is a potentially simple but yet powerful readout that allows distinguishing between healthy and diseased myoblast cells.

The characterization of nuclear deformations described above was conducted using classical image processing techniques. However, recent applications of generative deep learning models to various biological settings including microscopy image analysis^21^ and multi-omics data analysis^22^ motivated our interest in the potential applicability of such models to the study of nuclear deformations. Thus, the goal of the present study is to explore the capability of unsupervised deep learning algorithms to rapidly and automatically classify the different types of nuclear deformations on microgroove substrates and to investigate if such algorithms can enhance our ability to discriminate between normal and diseased cells. To this end, we have opted to use a variational autoencoder (VAE) architecture, which is one of the most popular approaches to generative modeling. VAE models capture an underlying data manifold, a lower dimensional space where data are distributed, and can disentangle sources of variation when trained on different classes of data. We assess the ability of a VAE to cluster 60×60 pixel visual patches, each containing a single nucleus, extracted and resized from 2048×2044 pixel images of myoblasts cultured on microgrooves. We show that this approach is effective in clustering nuclei based on their deformations with distinct clusters for myoblasts derived from healthy individuals and those derived from laminopathy patients.

## Results

### Diversity of nuclear deformations on microgrooves

Myoblast cells cultured on fibronectin-coated microgroove substrates (see schematic in Fig. 1a) and fixed a couple of hours after seeding exhibited various types of nuclear deformation with distinct nuclear morphologies. More specifically and as described in our previous work^20^, we were able to identify the following three categories of deformation corresponding to specific nuclear morphometric features examples of which are shown in the insets of Fig 2b: 1) “caged” nuclei that are fully confined or trapped within a groove and that are thus highly elongated and aligned in the groove direction (top inset), 2) “partly caged” nuclei that have a portion of their volume within a groove with the remainder suspended on an adjacent ridge (middle inset), and 3) “double caged” nuclei that have portions of their nuclei in two adjacent microgrooves with the remaining portion straddling the ridge separating the two grooves (bottom inset).

**Figure 1.**
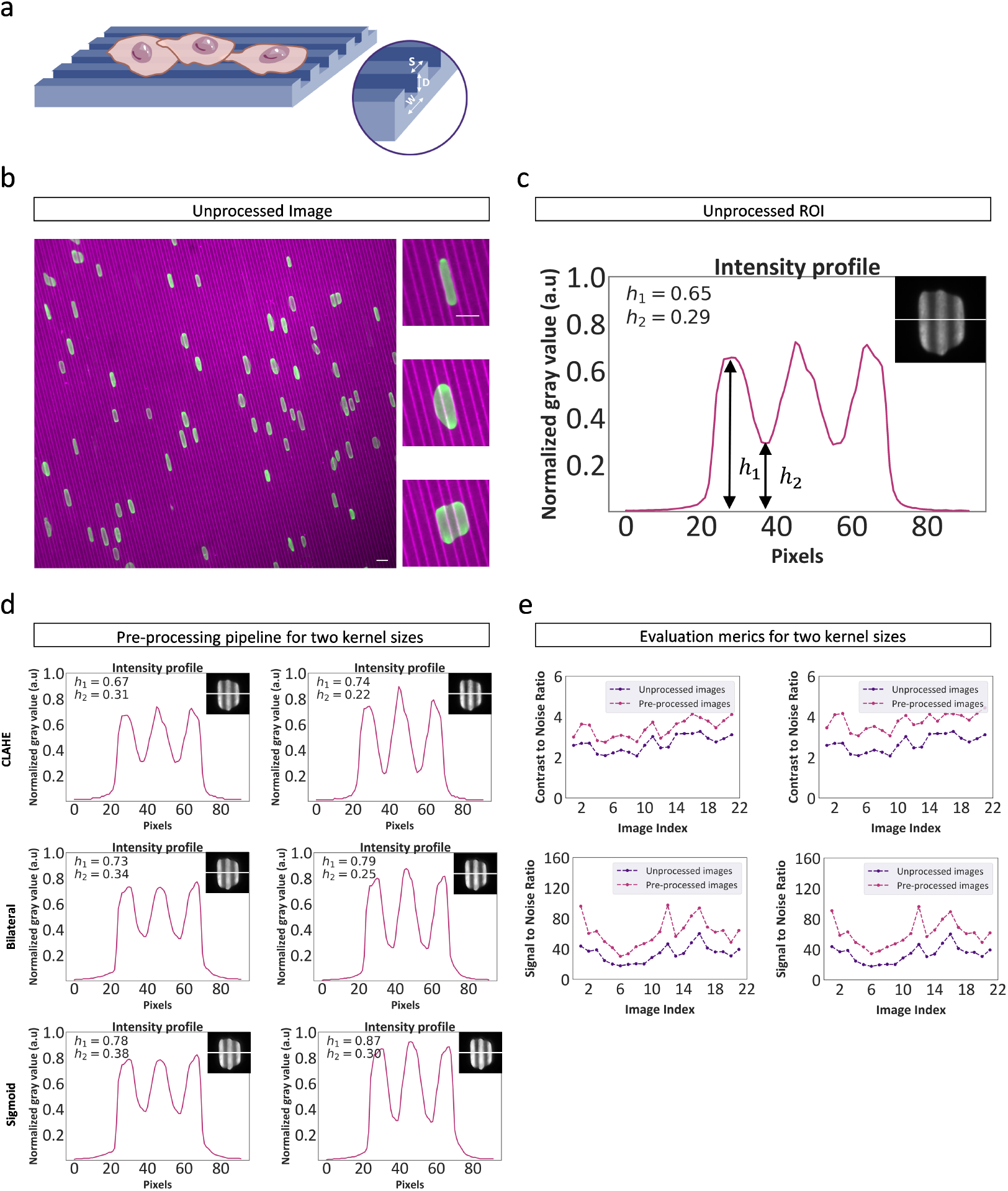
Effect of data processing with image enhancement. a) Schematic of myoblast cells cultured on fibronectin-coated microgroove substrates of spacing s, width w, and depth d. b) Immunostaining for fibronectin (grooves, magenta) and lamin-A/C (nucleus, green). Left: image acquisition with a 20X objective. Scale bar, 15*µm*. Right: Regions of interest (ROIs) for different configurations of nuclear deformations on the grooves. Scale bars, 15*µm*. c) Intensity profile along the line shown in the inset for the unprocessed image. The ROI contains a deformed nucleus on the groove surface. *h*_1_ and *h*_2_ are respectively the normalized intensity values for the peak and through one intensity peak corresponding to the parts of the nucleus lying on the groove and ridge areas of the microgroove substrate. d) Intensity profiles for the same conditions as in panel c following each step of the pre-processing pipeline for two different kernel sizes (left : 1*/*10× image width, right: 1*/*60× image width). e) Contrast-to-noise ratio (CNR) and signal-to-noise ratio (SNR) values with and without the pre-processing pipeline for the entire 21 image-dataset and for both kernel sizes.

**Figure 2.**
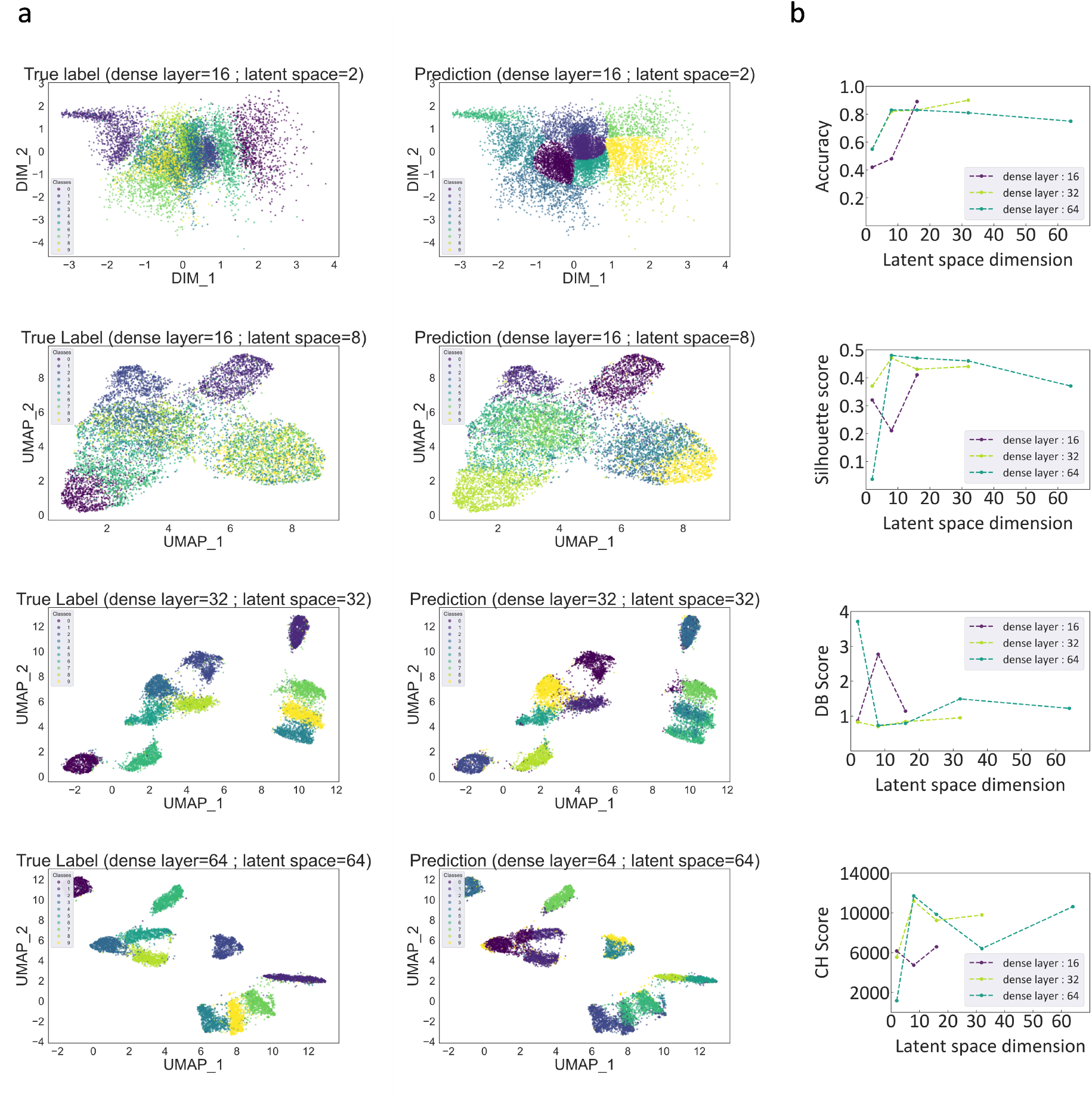
Calibration of our VAE for clustering the MNIST library. a) Latent space representation of true labels (left) and predicted labels (right) using a Gaussian Mixture Model (GMM) algorithm with a standard VAE, evaluated across different dense layer numbers and latent space dimensions. UMAP representation is used for latent space dimensions greater than 2. b) Evaluation of the clustering performance in the latent space with GMM for a standard VAE with different dense layer numbers (16, 32, 64) and latent space dimension (2, 8, 16, 32, 64). Scores chosen to evaluate the performance are: accuracy score, Silhouette score, Calinski-Harabasz (CH) Index, and Davies-Bouldin (DB) score. The accuracy score yields the fraction of correctly classified samples based on class equivalence between the predicted labels and the true labels. The Silhouette, Calinski-Harabasz, and Davies Bouldin scores are used to evaluate the quality of clustering of the predicted labels.

### Image contrast enhancement

Our complete dataset is comprised of 2048×2044 pixel grayscale images (see Figure 1b for an example) obtained from a number of independent experiments. Variability in signal intensity can therefore arise depending on immunofluorescence staining quality and/or microscope lighting conditions. When utilizing a VAE, images with low contrast or significant textural variations may introduce misleading features that influence our assessment and appreciation of nuclear deformations.

To evaluate the model’s sensitivity to these potentially problematic issues, we have devised an image pre-processing pipeline that consists of the following three steps: (1) contrast limited adaptive histogram equalization (CLAHE), (2) bilateral filtering, and (3) sigmoid correction. In the CLAHE step, the choice of clip limit and kernel size is paramount, as these parameters critically influence the quality of the enhanced image. Optimal values of these parameters were based on our assessment of their impact on both the contrast-to-noise ratio (CNR) and the signal-to-noise ratio (SNR) for a random dataset composed of 21 unprocessed and pre-processed images. This assessment revealed that setting the clip limit to 0.01 yielded the highest SNR and CNR values (Supplementary Figure S1a). As for kernel size, the CNR and SNR were found to remain relatively stable over a range of values, with a slight increase observed for kernels of approximately 30 pixels in size (Supplementary Figure S1b).

Building upon the findings above, we investigated the pipeline’s performance for two kernel sizes: approximately 30 pixels (1/60 of the image width) and 200 pixels (1/10 of the image width). This was accomplished by plotting the intensity profile along a line orthogonal to the cell major axis within a region of interest (ROI) that contains a nucleus deformed within the grooves (see inset of Fig. 1c). The intensity profiles along this line for both the unprocessed (Fig. 1c) and pre-processed (Fig. 1d) images show periodic peaks corresponding to the different parts of the nucleus positioned either on the groove or ridge surface. Application of the CLAHE algorithm (Fig. 1d, top panel) led to amplification of the intensity peaks for both kernel sizes, thus confirming the intended contrast enhancement. The fact that peak amplification was more pronounced for the smaller kernel size underscores the importance of carefully selecting this hyperparameter for the CLAHE algorithm. Subsequent addition of bilateral filtering (Fig. 1d, middle panel) increased the smoothness of the intensity profiles, resulting in less sharp intensity peaks and a reduction in textural variation. Finally, performing the sigmoid correction enhanced the contrast of the ROI (Fig. 1d, bottom panel).

At the level of the image of the nucleus, the final outcome of the pre-processing pipeline demonstrates an increase in signal clarity with reduced textural variation while preserving the edges of the nucleus. To quantify the effect of this pipeline for several ROIs, we evaluated the CNR and the SNR before and after pre-processing for the same 21-image dataset as in Fig. S1 and for the two kernel sizes considered. As depicted in Fig. 1e, the pipeline leads to increased CNR and SNR for both kernel sizes with slightly higher CNR values for the lower kernel size. We also evaluated the peak SNR (PSNR) as well as the structural similarity index measured (SSIM) for the same dataset (Supplementary Figure S2). Those results indicate higher quality of the pre-processed images while preserving similarities between the unprocessed and pre-processed images. Taken together, the results of this section underscore the added value of applying this pre-processing pipeline in the context of 2D images acquired via fluorescence microscopy on cells cultured on microgroove substrates.

### Calibration of our VAE for clustering the MNIST library

In this section, we aim to calibrate our VAE architecture by analyzing how the number of dense layers and the latent space dimensionality affect its performance in terms of clustering within the latent space embedding . To this end, we used the MNIST dataset, which consists of a large number of 28 × 28-pixel grayscale images of handwritten digits. While previous studies have commonly used the MNIST dataset to assess the clustering accuracy of novel models; there has been limited investigation into how varying the number of dense layers and the dimensionality of the latent space affect cluster representation and clustering performance—particularly in the context of applying a Gaussian Mixture Model (GMM), as we do here. We visualized the latent space embeddings of the true labels of the ten digits and their predictive clusters using a GMM algorithm as depicted in Figure 2a. For a dense layer number of 16 and a latent space dimension of 2, respectively, there is overlapping of the true labels, indicating that there is not a clear-cut distinction among the labels. The clusters become increasingly distinct as the number of dense layers and the latent space dimension are increased. We quantify these observations using various clustering metrics across different combinations of dense layer numbers and latent space sizes, as shown in Figure 2b. When building the VAE architecture, the latent space size was always less than or equal to the number of dense layers. When the sizes were equal, the encoder mapped the input data directly onto the latent space without any dimensionality reduction. The accuracy score ranges from 0.43 (for a dense layer number and latent space size of 16 and 2) to 0.9 (for a dense layer number and latent space size of 32 and 32). For a dense layer number of 64, the accuracy score decreases as the latent space size increases from 8 to 64 . The Silhouette, Calinski-Harabasz, and Davies-Bouldin scores exhibit similar trends as the accuracy score. When examining the accuracy score, the best result is obtained with a dense layer number and latent space size of 32 each. These results highlight the importance of hyper-parameter tuning for clustering assignment of the latent space when using a VAE. The current results also demonstrate the efficiency of a GMM algorithm for cluster assignment.

When evaluating various architectures on both the Fashion MNIST dataset—a more challenging alternative to the original MNIST—and our own training dataset, we consistently achieved the highest accuracy and silhouette scores with a dense layer number and latent space size of 32 each. Therefore, this architecture was selected for all subsequent clustering of nuclei on the microgroove substrates.

### Pre-processing tuning for clustering optimization on nuclei patches

Following calibration of the VAE, we now wish to tune our pre-processing to optimize the clustering performance on a training dataset of deformed myoblast nuclei on microgrooves. To this end, we investigated: 1) the impact of rescaling image patches with a ROI that meets the fixed dimension requirement of the VAE, and 2) the influence of the contrast enhancement pipeline on the latent space. The architecture of the VAE was maintained constant throughout the training. Three different cropping and rescaling sizes were tested: 28 × 28 pixels, 64 × 64 pixels, and 184 × 184 pixels. The 64 × 64 patch size, when trained with the VAE, show higher scores of the clustering performance metrics (Supplementary Figure S3). This value corresponds to the power of 2 closest to the median and average values of the maximum dimension of the bounding boxes extracted from our training dataset. We then evaluated the effect of the contrast enhancement on the clustering performance using the silhouette, Calinski-Harabasz, and Davies Bouldin scores. The best clustering scores were obtained for the CLAHE filtering with a kernel size of 1*/*60× image width, while the lowest scores were obtained for unprocessed images (Table 1). These results demonstrate that the clusters in the latent space are better separated with less overlap in the case of enhanced images (Figure 3). These results underscore the impact of image processing (both patch rescaling and contrast enhancement) for clustering visual features when using a VAE.

**Table 1.** Metrics used for cluster assignment (silhouette score, Davies Bouldin score, and Calinski-Harabasz index). The dataset used is composed of image patches of size 64×64-pixels and pre-processed with CLAHE with filter size of 1*/*10 × image width and 1*/*60 × image width.

|  | raw images | 1/10 | 1/60 |
| --- | --- | --- | --- |
| Silhouette Score | 0.17 | 0.2 | 0.31 |
| Davies Bouldin Score | 1.6 | 1.26 | 0.99 |
| Calinski-Harabasz Index | 8241 | 12 962 | 21 785 |

**Figure 3.**
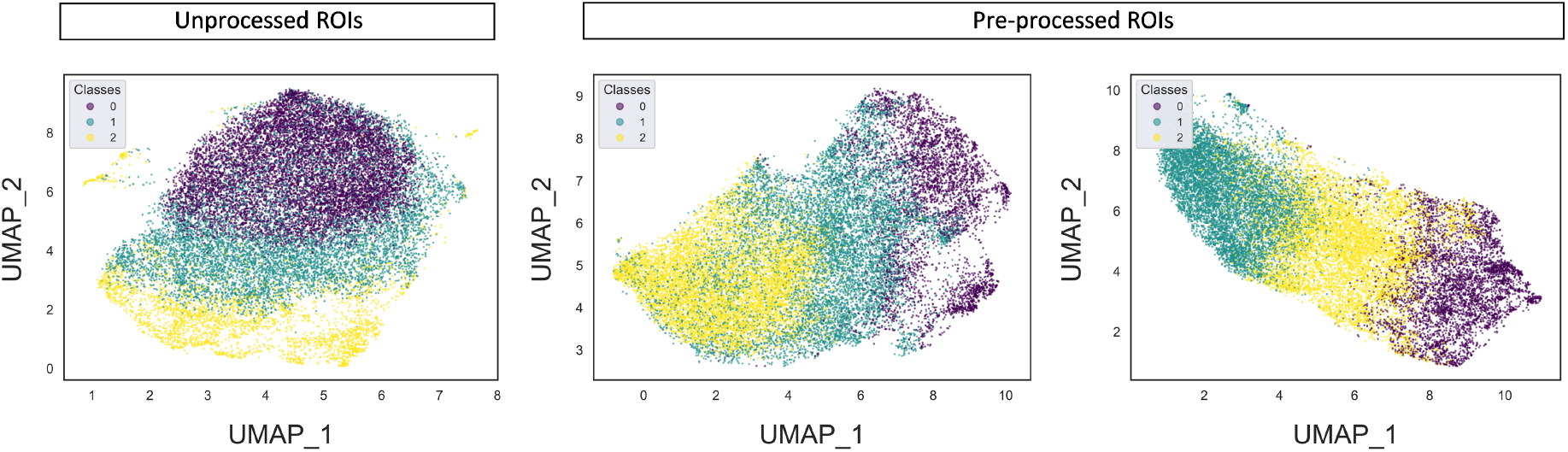
Pre-processing tuning for clustering optimization on nuclei patches. Visualization of the latent space embeddings for a training dataset with unprocessed images (left), pre-processed images using our pipeline with a kernel size of 1*/*10× image width (middle) and 1*/*60× image width (right). The number of clusters in the latent space is set to three. The different classes represent nuclei in different configurations from confined or trapped nuclei to deformed nuclei on multiple grooves.

### Clustering of morphological nuclei features using the optimized VAE

Now that the VAE parameters for a real dataset have been tuned to optimize clustering, we investigated its ability to segregate normal myoblasts from those derived from LMNA-related congenital muscular dystrophy patients. To this end, the deformations of myoblast nuclei within the microgrooves for both wild type (WT) cells and those carrying two different mutations, Mutant R249W and Mutant L380S, were studied. Compared to LMNA mutant nuclei, WT nuclei on microgrooves visually exhibit a greater tendency towards circular shapes with less morphological variability (Figure 4a).

**Figure 4.**
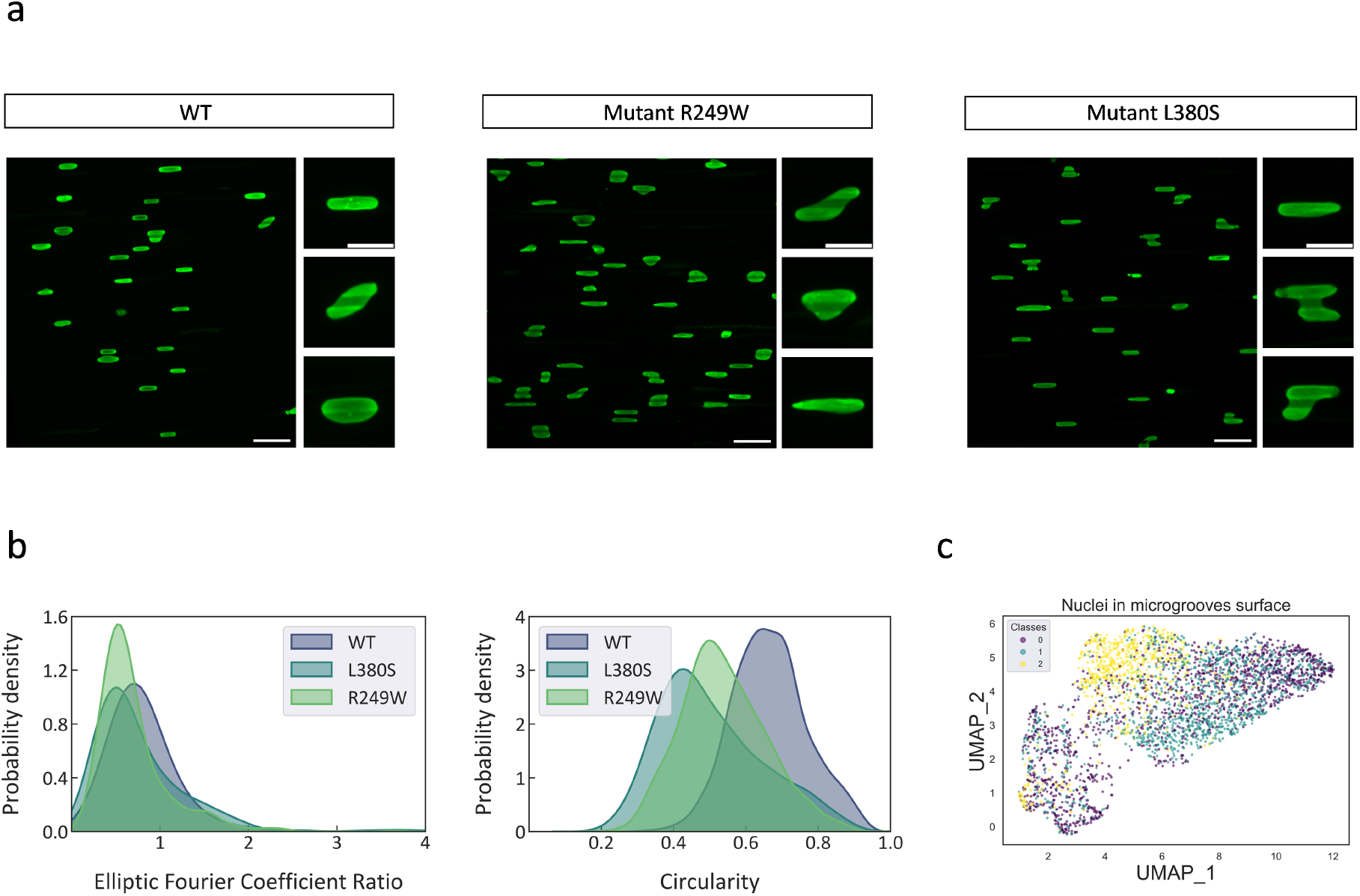
Clustering of morphological nuclei features using the optimized VAE. a) Immunostaining for lamin A/C (green) of WT and mutant myoblast cells on microgrooves. Insets show zoom-ions on particular cells to underscore morphological differences between WT and mutant cells. Scale bar for large images = 50 µm; scale bar for insets = 20 µm. b) Histograms of Elliptic Fourier Coefficient Ratio (EFCR) and circularity for WT and mutant myoblasts. c) Visualization of the latent space embeddings using UMAP representation for nuclei of the different myoblast cell types. Classes 0, 1, and 2 respectively represent R249W mutant cells, L380S mutant cells, and WT cells.

We tested two morphological metrics on ground-truth nuclei types: circularity and Elliptic Fourier Coefficient Ratio (EFCR). WT nuclei differ in circularity histograms and, to a lesser extent, in the EFCR (Figure 4b). Although these morphological metrics suggest a certain capability of discriminating between WT and mutant myoblasts (Figure 4b), there is significant overlap among the histograms. Applying our VAE to nuclei patches of the different cell types and looking at the latent space (Figure 4c) reveals two different clusters represented by WT and mutant cells; however, no clear demarcation between the latent space representations of the two mutants is evident.

We had previously shown that nuclear deformations on microgrooves can assume several configurations including complete confinement or “caging” within the grooves, “partial caging” where a part of the nucleus is in a groove and the rest on a ridge, and “double caging” where the nucleus straddles a ridge and has portions of it that penetrate two adjacent grooves^20^. We wished to establish if our VAE was capable of discriminating among these different deformation configurations or of even potentially identifying additional more subtle deformation categories for both WT and mutant myoblasts. To this end, we examined the latent space for WT and mutant myoblasts assuming the presence of 3, 4, or 5 clusters (Fig. 5a). When the number of clusters was set to 3, the clusters were quite distinct for both the wild type (WT) and the two types of mutant myoblasts but we observed notable differences in clustering scores between the wild type and the L380S mutation. Increasing the number of clusters to 4 or 5 did not significantly improve clustering results (Fig. 5b). However, when evaluating the UMAP projection, we observed more cluster overlap for the wild type compared to the two mutant lines. Specifically, the L380S mutation exhibited five easily distinguishable clusters, whereas the wild type did not. These findings suggest that the three categories of nuclear deformations as described in previous work^20^ may be insufficient to capture the full range of deformations observed on microgroove substrates, especially in the case of mutated cell lines.

**Figure 5.**
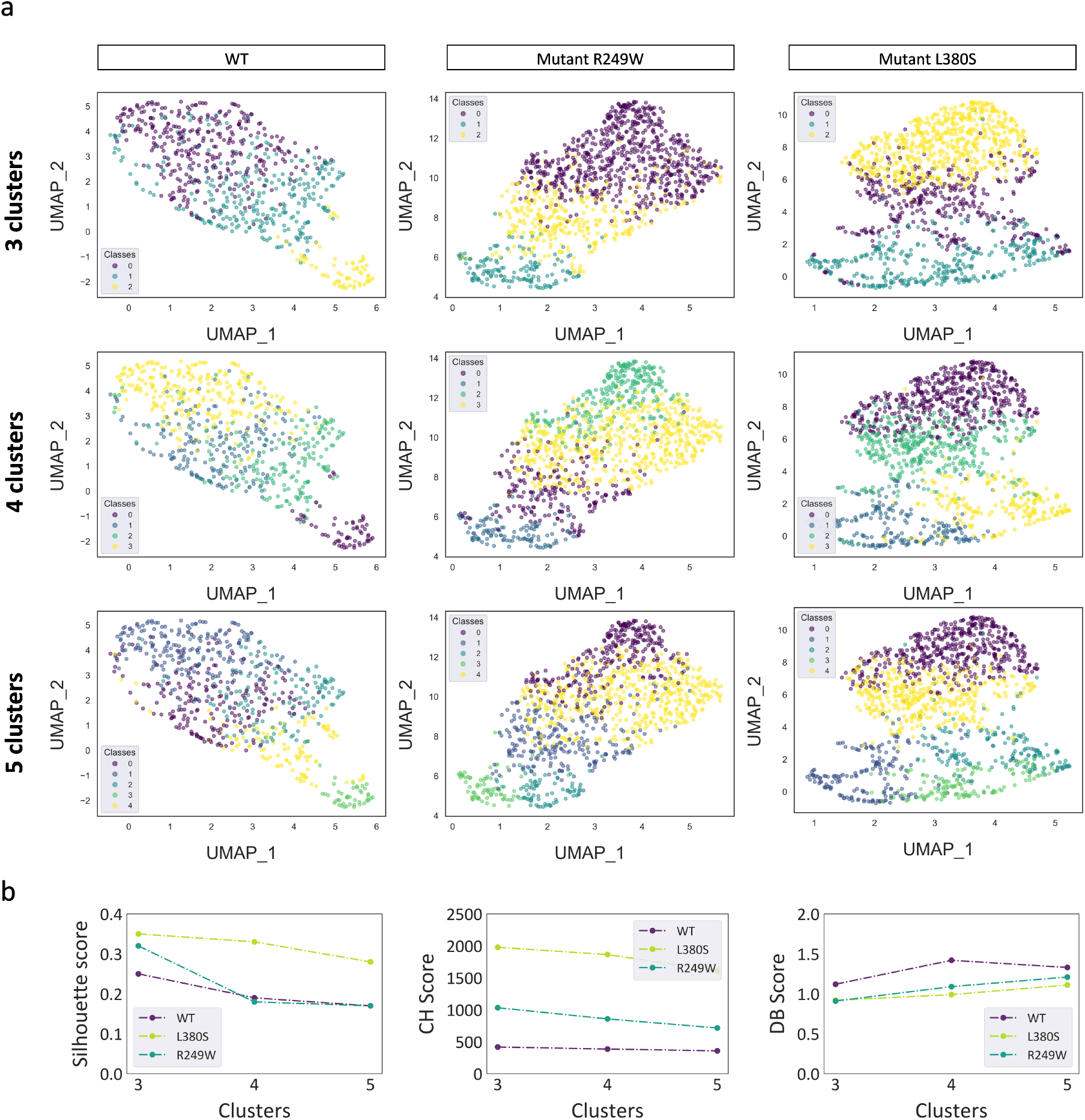
Clustering of morphological features using the optimized VAE. a) Visualization of the latent space embeddings of respectively WT cells, L380S mutant cells, and R249W mutant cells using UMAP representation after clustering. The number of clusters is set to 3, 4, or 5. The different classes represent different degrees of deformation of the nuclei on the grooves. b) Quantitative assessment of clustering performance for the different cell types and for the different number of clusters (3, 4, or 5) using the Silhouette score, the Calinski-Harabasz index, and the Davies Bouldin score.

By categorizing each nucleus within the clusters derived from latent space embeddings, we can calculate the percentage of nuclei in different deformation categories for both wild type and mutant myoblasts. The number of clusters chosen for this categorization was obtained through visual inspection of clearly distinct clusters in the UMAP representation. Interestingly, while this visual inspection revealed three clusters for WT myoblasts, four clusters were observed for the mutant cells. This difference had its origin principally in the double caged category. More specifically, while double caged nuclei for WT cells were broadly oval in shape, those in mutant cells were either H-shaped with symmetric caging ion either side of a ridge or asymmetric caging with the two caged portions having significantly different sizes (see Fig 6a).

**Figure 6.**
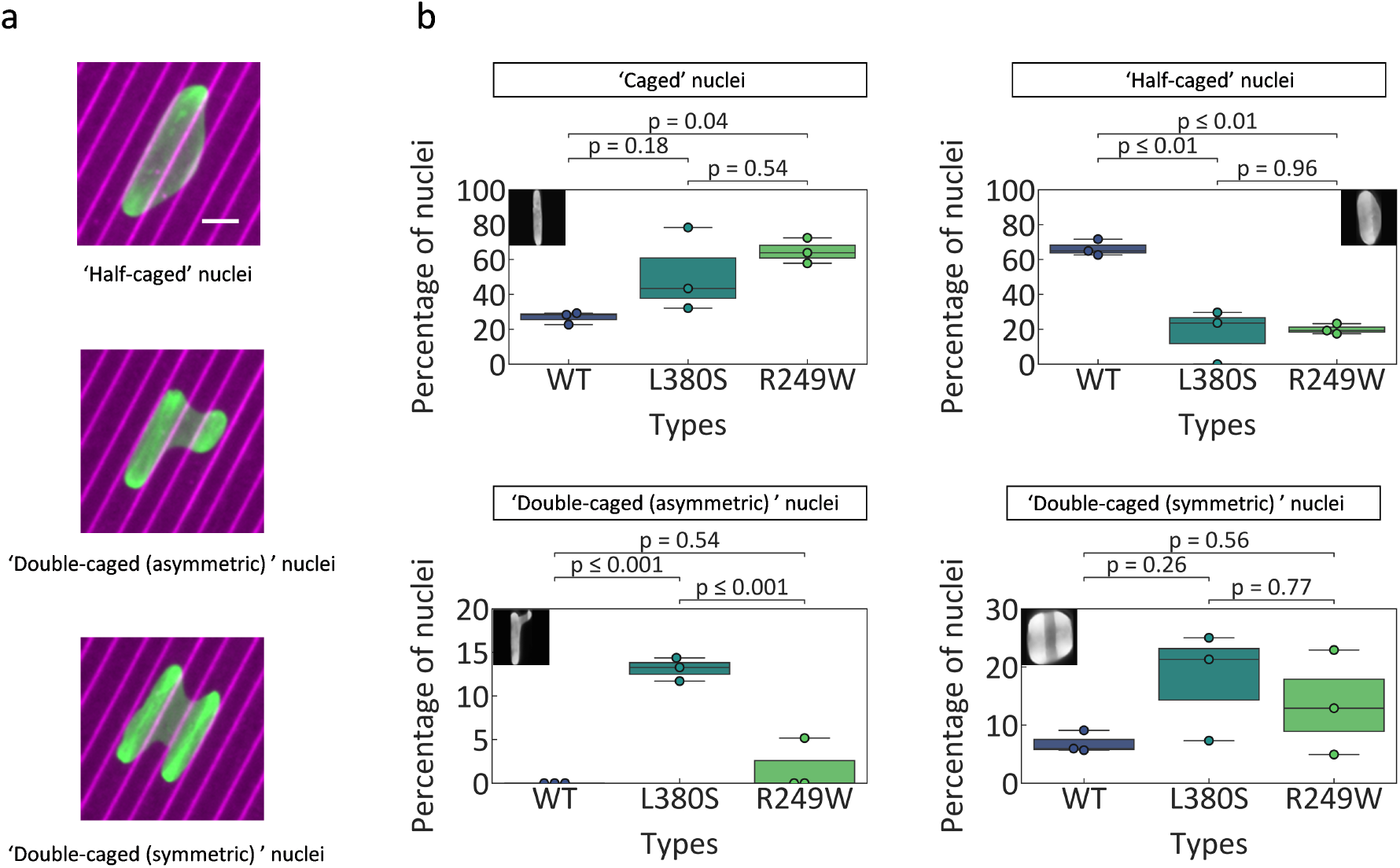
Clustering of morphological features using the optimized VAE. a) Different configurations of mutant myoblasts: the top panel corresponds to half-caged nuclei, the middle panel to double-caged nuclei (asymmetric), and the bottom panel to double-caged nuclei (symmetric). Image acquisition with a 20X objective. Scale bar, 15*µm* b) Percentage of nuclei within each deformation category for WT, L380S, and R249W mutant myoblasts. The number of nuclei for each group has been computed using our optimized VAE and a GMM algorithm to cluster the embeddings in the latent space. The number of nuclei patches per experiment varies between 200 and 500.

As shown in Fig. 6, WT myoblasts exhibited considerably less full nuclear caging, and consequently considerably more partial caging, than mutant myoblasts.

These results demonstrate that the use of our VAE for clustering of various categories of nuclear deformations on microgroove substrates has some potential as an effective approach for distinguishing between healthy and diseased myoblast cells.

## Discussion

In this study, we explored the potential of a standard VAE to classify different types of nuclear deformations on microscopy image patches from myoblasts cultured on microgroove substrates that mimic the anisotropic topography of the basement membrane and that have been shown to induce significant nuclear deformations in multiple cells types^20^. Of note is that we integrated a GMM clustering algorithm into the latent space, enhancing our ability to capture intricate data distributions. To perform the desired classification, we developed a pre-processing pipeline that enhances the visibility of lighting variations, which accentuates nuclei deformation on the microgrooves, while also mitigating the impact of intensity variations caused by different experimental conditions. We calibrated our VAE with a GMM using the MNIST library and showed the effect of this pre-processing pipeline on clustering performance using different clustering metrics including the Silhouette, Davies Bouldin, and Calinski-Harabasz scores. This analysis demonstrated the importance of the pre-processing pipeline in improving the clustering performance.

In our training set, distinct clusters were established corresponding to varying degrees of nuclear deformations including nuclei that were fully ‘caged’ within the grooves and nuclei that were ‘deformed’ within two adjacent grooves. Because the extent of nuclear deformations depends on nuclear mechanical properties, it would be expected that pathologies characterized by abnormalities in nuclear mechanics such as laminopathies would lead to nuclear deformations on microgrooves that differ from those of normal cells. To investigate the ability of our VAE to capture such differences, we tested a data set of nuclei from both control myoblast cells and myoblast cells with two different mutations from patients with LMNA-related congenital muscular distrophy. In parallel, we also evaluated several morphometric features in these three different cell phenotypes including circularity and the Elliptic Fourier Coefficient Ratio. Interestingly, basing the comparisons on these morphometric parameters revealed that circularity provides superior discriminatory capability among the three cell phenotypes than the elliptic Fourier coefficient ratio. This underscores a principal limitation of clustering based on morphometric features, which is the inability to know a priori which morphometric parameters are the most pertinent ones for a particular cell type. This notion has been highlighted in several previous studies. For instance, Prulovic et al. explored the ability to discriminate among cells derived from tumors of different grades based on a broad array of cellular and nuclear morphometric parameters and reported that area, convexity, and circumference were the most pertinent^23^. Antmen et al. investigated nuclear deformations on micropatterned surfaces in different types of breast cancer cells with the goal of distinguishing among their malignancies and determined that significant differences were only evident for certain morphometric parameters but not others^24^.

Employing deep learning techniques promises to provide a more automated approach to characterizing distinct cellular morphologies, potentially driven by visual patterns not captured by a single morphometric measure. Some studies have used deep learning to characterize nuclear morphology in various settings. For instance, Heckenbach et al. used nuclear morphology as the basis for predicting human fibroblast senescence with an accuracy of up to 95%^25^. To our knowledge, there have been no studies to date using deep learning techniques to assess nuclear deformations, particularly in the context of diseases such as laminopathies. While a limited number of studies have explored morphometric features of nuclei in laminopathies, the assessment of nuclear deformations is typically qualitative^26^. Employing a standard VAE with GMM algorithm as demonstrated here permits unsupervised discovery of nuclei deformation phenotypes and automated classification of nuclear deformations. The resulting clustering allows discrimination between healthy (wildtype) cells and those carrying laminopathy-associated mutations.

A limitation of employing a VAE with GMM clustering algorithm is that the determination of the number of clusters relies on manual selection through visual analysis of various modes of deformation of nuclei on the microgrooves. This necessitates human oversight and involvement. To address this issue, integrating the GMM clustering into the learning process would obviate the need for predefining the number of clusters^27^. Another drawback of such unsupervised neural networks is that the output is simply a group of clusters without information as to what these clusters correspond, which limits the applicability to direct medical applications. Using supervised classification in place of unsupervised clustering promises to yield more medically relevant labels. In addition, incorporating established methods such as guided backpropagation^28^ or Grad-CAMs^29^ would provide insight into visual features that drive the learning representation.

In conclusion, our study presents a novel approach for discovering and classifying nuclear deformations on micropatterned substrates, with special focus on laminopathies. We propose an unsupervised deep learning method utilizing a VAE. This approach allows discrimination between healthy cells and those carrying laminopathy-associated mutations based on nuclear deformation characteristics, marking the first instance of such an approach. These findings pave the way for integrating deep learning with the study of nuclear deformation and mechanics, offering new avenues of exploration in this field.

## Methods

### Fabrication of microgrooved substrates

The original microstructured silicon wafer was fabricated using laser lithography (Heidelberg µPG101) of a layer of SU8-2010 (MicroChem, USA). After exposure to trichloro(1H,1H,2H,2H-perfluorooctyl)silane (Sigma) vapor for 20min, the silicon wafer was used to create PDMS (Sylgard 184, Sigma Aldrich, 1:10 ratio) replicates. To create the final coverslip on which the cells were cultured, liquid PDMS was spin-coated at 1500 rpm for 30 s on the PDMS mold. Before reticulation overnight at 70 °C, a glass coverslip was placed on top of the PDMS layer. After reticulation, the glass coverslip attached to the microstructured PDMS layer was gently demolded with a scalpel and isopropanol to facilitate detachment. Microstructured coverslips were then sonicated for 10 min in ethanol for cleaning and finally rinsed with water. The final structure used for the experiments is composed of PDMS 5 µm width and 4 µm deep microgrooves that are apart by 5 µm.

### Cell culture and image acquisition

Study was performed on immortalized human myoblasts from control subject without muscular disorders (WT) and patients carrying the following heterozygous LMNA mutations responsible for severe congenital disorders^30^ : LMNA p.Arg249Trp (hereafter referred to as R249W) and LMNA p.Leu380Ser (hereafter referred to as L380S). Cells were cultured on the microgroove surfaces for 24 h. after which the cells were fixed with 4 % paraformaldehyde (Thermo Fisher) in PBS for 15 min. After 1 h in a blocking solution containing 0.25% Triton and 2% bovine serum albumin (BSA), cultures were incubated for 1 h at room temperature with primary antibodies. All antibodies were diluted 1/400-1/200 in a solution containing 0.25% Triton and 1% BSA. Coverslips were washed three times with PBS and incubated for 1 h at room temperature with Alexa Fluor 555-conjugated donkey anti-rabbit antibody (ab150074, Abcam) or Alexa Fluor 488-conjugated donkey anti-mouse antibody (ab150105, Abcam) as well as DAPI for nuclear detection. Epifluorescence images were acquired on an inverted microscope (NikonEclipse Ti) with a 20 X objective (Nikon Plan Fluor NA = 0.5).

### Images processing

A pre-processing pipeline was developed for contrast enhancement, noise reduction and edges preservation. The pipeline consists of the following steps of spatial domain transformation which were implemented using the scikit-image librairy :

- CLAHE (contrast limited adaptative histogram equalizer) method is based on an adaptive histogram equalizer where the input image is subdivided into non-overlapping blocks (tiles or kernels). This methods enhances the contrast within each of the blocks individually. A ‘clip-limit’ controls the degree of contrast enhancement by setting each histogram to a predefined value and redistributing the clipped histogram equally across the histogram. In the present work, the clip limit is fixed at 0.01 based on an evaluation of its effect on various performance metrics. The pipeline is tested for two kernel sizes: 1*/*10 × image width and 1*/*60 × image width.
- A bilateral filter (BF) is an edge preserving spatial and noise reducing filter. It is formulated as follows :

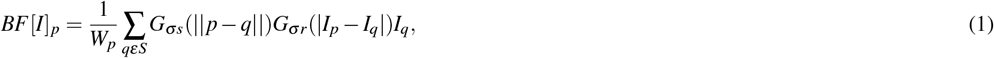

where *W*_*p*_ is a normalization factor, *G*_*σs*_ a spatial Gaussian that decreases the influence of distant pixels, *G*_*σr*_ a range Gaussian that decreases the influence of pixels q with an intensity value different from *I*_*p*_. We used the restoration.denoise_ bilateral function in scikit-image with the default parameters.
- A sigmoidal correction for contrast adjustment was applied. This function, implemented in Python, transforms the input image pixelwise so that the relationship between the output (0) and the input (I) is as follows :

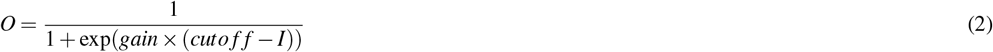

after scaling each pixel to the range 0 to 1. The gain was fixed at 3 and the cut-off at 0.2.

Images of microgrooves and nuclei were oriented in the same direction to avoid orientation learning by the VAE. The nuclei were segmented using the Li-thresholding method. The unprocessed images and coordinates of the bounding box were used to crop unprocessed and pre-processed images into several images containing a single region of interest (ROI). Cropped images were then rescaled to fixed sizes (28 × 28; 64 × 64; 128 × 128). The aspect ratio was preserved using padding techniques. The pixel value used for padding was equal to the minimum pixel value of the cropped image. To increase our training dataset, we flipped the images in the up / down / left / right directions. The training dataset was composed of 21 732 images of nuclei deformed in the grooves.

### Performance metrics for pre-processing

For the purpose of this study, the performance metrics chosen to extract certain features from our pre-processed images included the contrast-to-noise ratio (CNR), signal-to-noise ratio (SNR), peak signal-to-noise ratio (PSNR), and the Similarity Index (SI). We assessed these metrics for a dataset of 21 images acquired during different experiments.

The CNR metric assesses the ability of the pre-processing pipelines to differentiate between two regions with different backscattering intensity. In this study, we defined the CNR as follows :

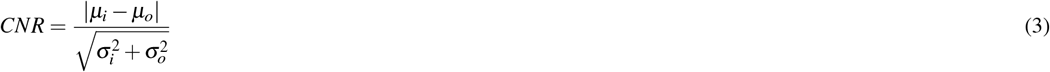

where *µ*_*i*_ and *µ*_*o*_ are the mean pixel values inside and outside our ROI and *σ*_*i*_ and *σ*_*o*_ are respectively the variance of the signal power inside and outside the ROI.

The SNR is defined as the power ratio between a signal and the background noise and is computed as :

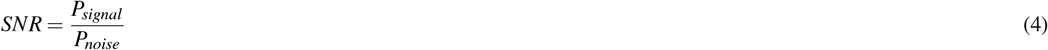

where *P*_*signal*_ is the average power of the pixels within the ROIs and *P*_*noise*_ is the standard deviation of the pixels within the regions that contains only noise.

The PSNR and SI were implemented using the scikit-image Python librairy. PSNR, defined as the ratio between the maximum possible power and the corrupting noise that affects image representation, is defined as follows :

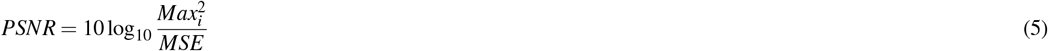

where Max is the maximum possible pixel value of the image and MSE is the Mean Squared Error and represents the average squared difference between the reference signal and the reconstructed signal.

The SI, which ranges from 0 to 1, is defined as the fraction of pixels in the enhanced image that matches the pixels in the primary or unprocessed image.

### VAE model

A VAE learns probabilistic representations z given input x and uses these representations to reconstruct the input x’. The generative model learns a joint distribution *p*_*θ*_ (*x, z*), with a prior distribution over latent space *p*_*θ*_ (*z*), and a stochastic decoder *p*_*θ*_ (*x*|*z*). A stochastic encoder *q*_*ϕ*_ (*z*|*x*), also called inference model, approximates the true but intractable posterior *p*_*θ*_ (*z*|*x*) of the generative model.

To approximate the true posterior, we need to minimize the Kullback-Leibler (KL) divergence between the introduced posterior and the true posterior,

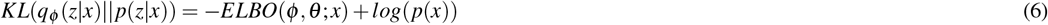

where *ELBO*(*ϕ, θ* ; *x*) is the evidence lower bound on the log probability of the data because the KL term must me greater than or equal to zero. Because the KL term is intractable to compute, minimizing the KL divergence is equivalent to maximizing the lower bound which can be written as follows :

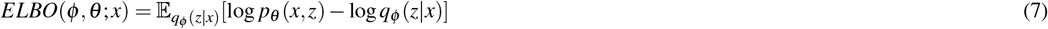

A standard VAE developed in Keras was used for the purpose of this study. Both encoder and decoder blocks consist of three layers, and all layers utilize a Rectified Linear Unit activation except the final output, which uses sigmoidal activation. All models were trained for 20 epochs (determined by identifying the loss function plateau), with batch size of 32. The shape of the dense layer and the latent space were adjusted using the MNIST and the fashion MNIST librairies and maintained constant for hyperparameter tuning of the training dataset.

Standard VAEs utilized the standard Evidence Lower Bound loss format characterized by reconstruction and divergence terms. We used a Binary Cross-Entropy (BCE) loss as the reconstruction term. The standard VAE loss used in this study can be defined as :

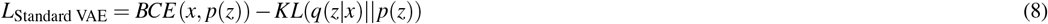

We applied the adaptive moment estimation (Adam) optimizer for gradient descent with default parameters (learning rate: 0.001, epsilon: 1e-07).

### Latent space clustering using a Gaussian Mixture Model

A Gaussian mixture model (GMM) is a probabilistic model where data samples are assumed to be generated by a mixture of *K* Gaussian distributions, whose parameters are estimated by an iterative model known as Expectation-Maximization (EM algorithm) for finding a local maximum likelihood. The Gaussian distribution of a vector *X* ∈ ℝ^*D*^ is characterized by its mean *µ* and its covariance matrix Σ :

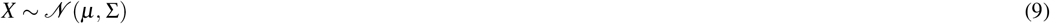

The Expectation-Maximization algorithm is based on generative models with *z* the vector of latent variables (*z*_1_, …, *z*_*n*_). GMM is the marginal distribution of the joint distribution :

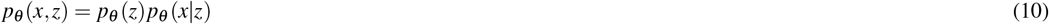

with :

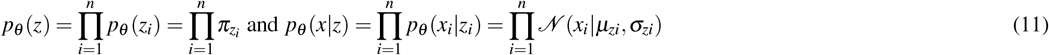

*z*_*i*_ ∈ 1, …, *K* and 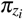 is the mixture proportion representing the probability that *x*_*i*_ belongs to the k-th mixture component. Given the latent variables, the log-likelihood is easier to compute and becomes :

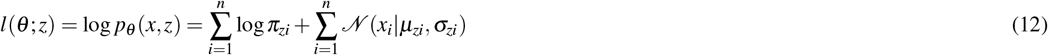

In order to evaluate the model’s ability to separate labeled cell populations, GMM clustering was applied to the encoding space using sklearn.

### Evaluation metrics

UMAP embeddings were calculated using the UMAP Python package. Clustering performance was evaluated using the accuracy score, the Silhouette score, the Davies Bouldin score and the Calinski-Harabasz index. The

Silhouette score measures the relation between cluster cohesion and cluster separation. It is calculated using the mean intracluster distance (*a*) and the mean nearest-cluster distance (*b*) for each sample. The Silhouette Coefficient for a sample is (*b*− *a*)*/max*(*a, b*). The Silhouette score ranges from -1 to 1, where positive values indicate a high separation between clusters, negative ones indicate that a sample has been assigned to the wrong cluster and a score of zero indicates that the data are uniformly distributed throughout the Euclidean space.

The Davies Bouldin Index is a measure of similarity between clusters. It is defined as :

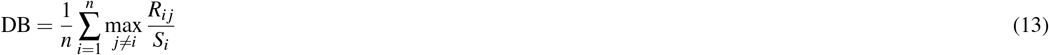

Where 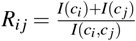, *I*(*c*_*i*_) being the mean distance between samples from cluster *c*_*i*_ and the center of the cluster, and *I*(*c*_*i*_, *c* _*j*_) the distance between the center of clusters *c*_*i*_ and *c* _*j*_. Lower values indicate better clustering.

The Calinski-Harabasz score also known as the Variance Ratio Criterion is defined as the ratio of the sum of between-cluster dispersion and of within-cluster dispersion. A higher score indicates better defined clustering. It is defined as follows :

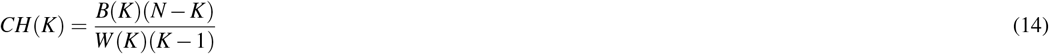

where K is the corresponding number of clusters, B(K) is the inter-cluster divergence, also called the inter-cluster covariance, W(K) is the intra-cluster covariance, and N is the number of samples. Larger B(K) values indicate a higher degree of dispersion between clusters. Smaller W(K) values point to a closer relationship in the cluster. A higher ratio, and thus a larger CH index, denotes better clustering.

### Statistical Analysis

All biological analyses were based on three independant experiments. Statistical analyses were performed using the scipy librairy implemented in Python. Multiple groups with a normal distribution were compared using a one-way ANOVA followed by Tukey’s posthoc test. The data points for each experiment and the significance levels are noted in the corresponding figure.

