## Supplementary information for "A Variational Autoencoder Model for Clustering of Cell Nuclei on Microgroove Substrates: Potential for Disease Diagnosis"

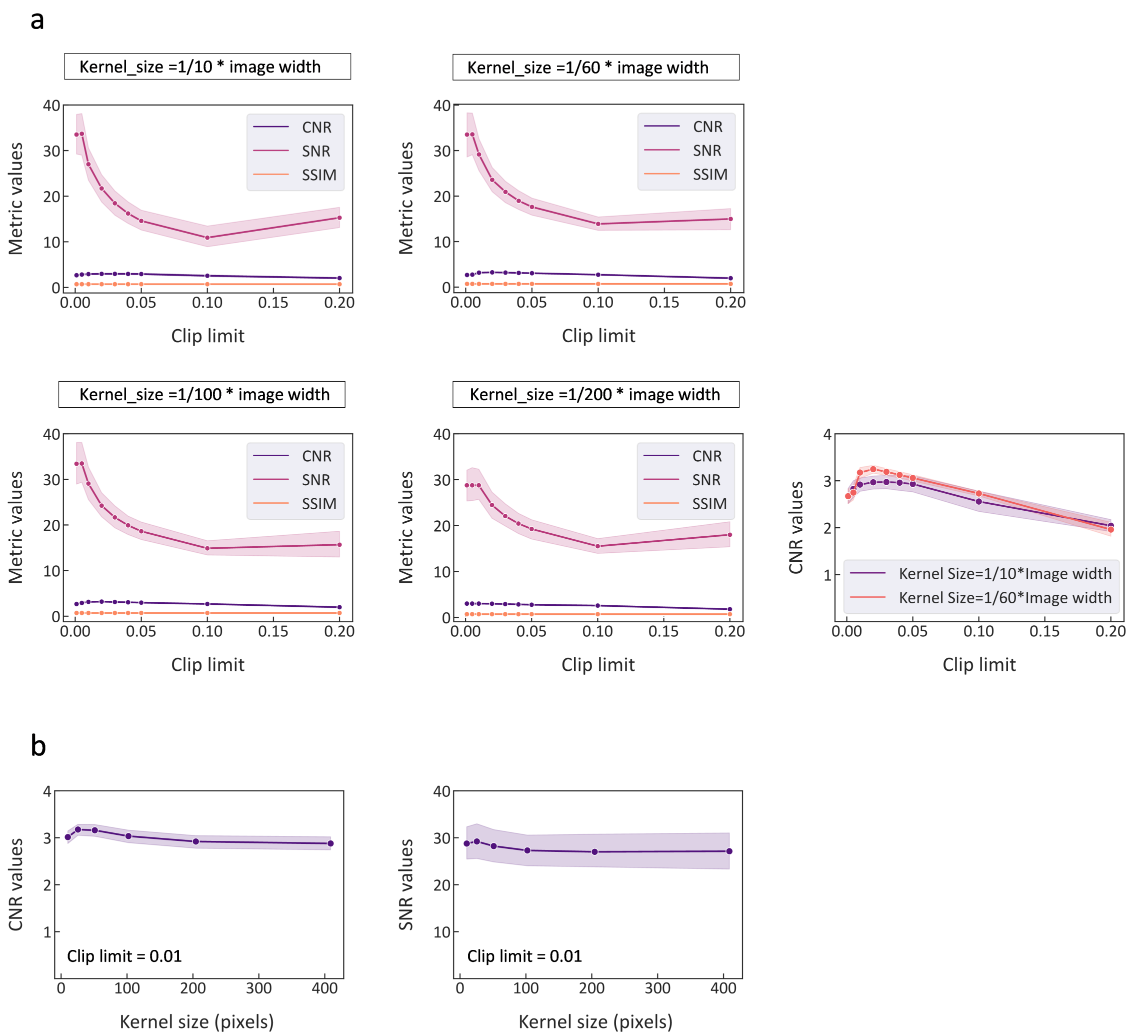


**Figure S1.** a) Evaluation of Contrast-to-Noise Ratio (CNR), Signal-to-Noise Ratio (SNR), and Structural Similarity Index (SSIM) as functions of clip limit for CLAHE-processed images with four distinct kernel sizes. b) Evaluation of Contrast-to-Noise Ratio (CNR) and Signal-to-Noise Ratio (SNR) for CLAHE-processed images with various kernel sizes, using a fixed clip limit of 0.001.

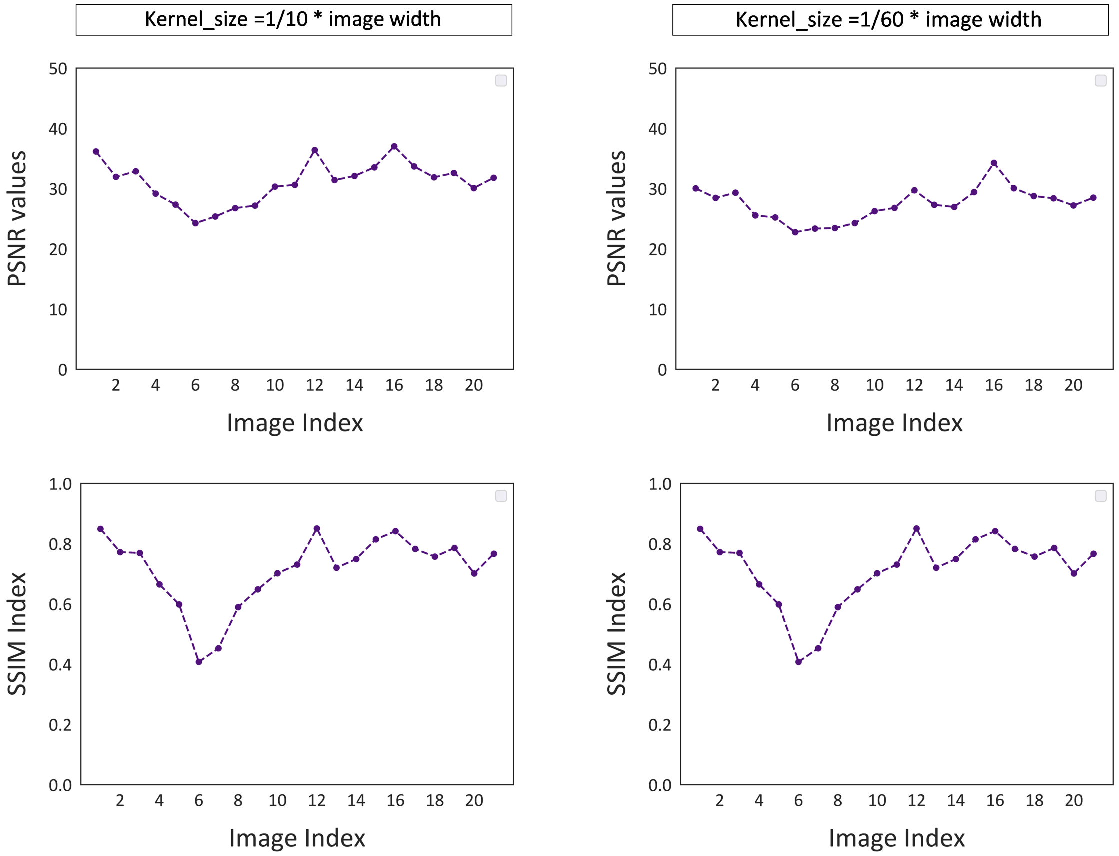


**Figure S2.** Assessment of Peak Signal-to-Noise Ratio (PSNR) and Structural Similarity Index (SSIM) for CLAHE-processed images, with two different kernel sizes : 1/10 × image width (left) and 1/60 × image width (right).

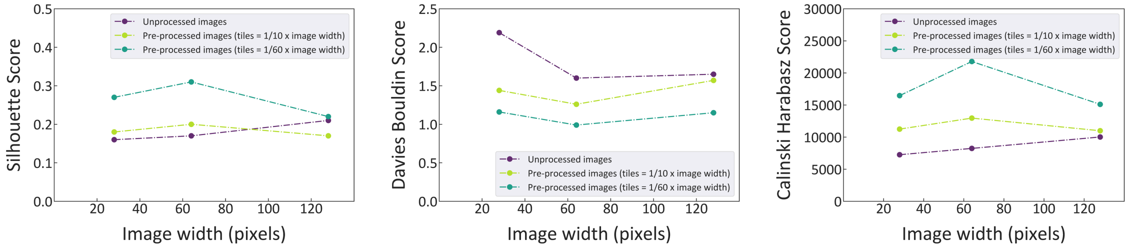


**Figure S3.** Evaluation of the Silhouette score, Davies-Bouldin score, and Calinski-Harabasz index for various nuclei patch sizes in both unprocessed and CLAHE-processed images, using two distinct kernel sizes : 1/10 × image width and 1/60 × image width.
